# High robustness of cytosolic glutathione redox potential under combined salt and osmotic stress in barley as revealed by the biosensor Grx1-roGFP2

**DOI:** 10.1101/2022.12.22.521445

**Authors:** Finja Bohle, Alina Klaus, Hendrik Tegethof, Markus Schwarzländer, Frank Hochholdinger, Andreas J. Meyer, Ivan F. Acosta, Stefanie J. Müller-Schüssele

## Abstract

Barley is a staple crop of major global importance and relatively resilient to a wide range of stress factors in the field. Transgenic reporter lines to investigate physiological parameters during stress treatments remain scarce.
We generated and characterized stable homozygous barley lines (cv. Golden Promise Fast) expressing the genetically-encoded biosensor Grx1-roGFP2, which indicates the redox potential of the major antioxidant glutathione in the cytosol.
Our results demonstrate functionality of the sensor in living barley plants. We determined the glutathione redox potential (*E*_GSH_) of the cytosol to be in the range of −308 to −320 mV. *E*_GSH_ was robust against a combined NaCl (150 mM) and water deficit treatment (−0.8 MPa) that caused growth retardation and showed only a minor oxidation after 96 h of treatment.
We conclude that the generated reporter lines are a novel resource to study stress resilience in barley.

**One sentence summary:** Generation and characterization of barley plants stably expressing Grx1-roGFP2 reveal high robustness of cytosolic glutathione redox potential (*E*_GSH_) under combined salt and osmotic stress.

## INTRODUCTION

Barley is among the most important crop species worldwide (Newton *et al*. 2011) and exhibits a high abiotic stress tolerance as compared to other cereal crops (FAO, https://www.fao.org/3/Y4263E/y4263e0e.htm; Munns *et al*. 2006). Varieties, landraces and wild barley accessions form a valuable genetic reservoir to screen for factors influencing resilience to the extreme conditions associated to climate change (Dawson *et al*. 2015; Muzammil *et al*. 2018). Several markers affecting winter survival, flowering time and abiotic stress tolerance have already been identified and are used in breeding (Newton *et al*. 2011; Dawson *et al*. 2015; Muzammil *et al*. 2018). Transcriptomic datasets investigating the response of barley to salt and/or water deficit identified oxidation-reduction processes as part of the stress response, especially after several days of stress exposure (Kreszies *et al*. 2019; Osthoff *et al*. 2019).

With its diploid, sequenced genome (Mascher *et al*. 2017), barley is a prime genetic model for the cereal grasses of the Triticeae tribe. Moreover, the variety Golden Promise is genetically accessible via embryo transformation (Imani *et al*. 2011; Amanda *et al*. 2022). Drought stress affects plant height, developmental timing and spikelet development with a role of PHOTOPERIOD-H1 (Ppd-H1), as demonstrated via introgression lines (Gol *et al*. 2021). However, while transgenic reporter lines are a standard research tool in other model plants, only few fluorescent reporter lines have been generated to gain insight into barley growth and physiology, as e.g. for auxin and cytokinin signalling (Kirschner *et al*. 2018; Amanda *et al*. 2022) or calcium signatures (Giridhar *et al*. 2022).

Genetically encoded biosensors are fluorescent proteins (or protein pairs) that change their properties specifically in response to a physiological parameter. As any other protein, they can be targeted to specific subcellular localizations with targeting signals. The development and characterization of biosensors for a wide array of physiological parameters has paved the way for the specific assessment of physiological states of subcellular compartments, ultimately challenging and shifting many paradigms of cell biology (reviewed in Meyer *et al*. 2021; Müller-Schüssele, Schwarzländer, *et al*. 2021). For example, the redox potential of the redox pair of glutathione (GSH) and its oxidized form glutathione disulfide (GSSG) has been revealed to be far from its midpoint potential in subcellular compartments harboring a glutathione reductase (GR), with only nanomolar amounts of GSSG present. This results in glutathione redox potentials (*E*_GSH_) as low as c. −320 mV in the cytosol of plant cells, making the largely reduced pool of glutathione a versatile electron donor reservoir for enzymes that draw electrons from GSH to perform reduction reactions. This includes enzymes involved in the scavenging of reactive oxygen species (ROS) or oxidative damage repair, e.g. dehydroascorbate reductase (DHAR) or atypical methionine sulfoxide reductases B (MSRB1) (Foyer & Noctor 2011; Rey & Tarrago 2018; Müller-Schüssele, Bohle, *et al*. 2021). The compartment-specific study of *E*_GSH_ has been largely driven by the characterization of redox-sensitive GFP2 (Meyer *et al*. 2007; Schwarzländer *et al*. 2008) that contains a GSH-dependent thiol switch on the outer face of the GFP beta-barrel structure. The redox status of this cysteine pair is dependent on *E*_GSH_ while reaction kinetics are accelerated by class I glutaredoxin (GRX) catalysis, resulting in roGFP2 linked to human glutaredoxin 1 (Grx1-roGFP2) as the most widely used *E*_GSH_ biosensor (Gutscher *et al*. 2008; Schwarzländer *et al*. 2016; Müller-Schüssele, Schwarzländer, *et al*. 2021).

The study of redox networks in *Arabidopsis thaliana* has already revealed that glutathione-dependent and thioredoxin-dependent systems can at least partially compensate for each other in mitochondria and the cytosol (Marty *et al*. 2009, 2019). This partially redundant input of electrons via glutathione reductase or the NADPH-dependent thioredoxin reductase A and B (NTRA, NTRB) largely prohibits oxidation of the glutathione pool. Nevertheless, null mutants of the cytosolic/peroxisomal glutathione reductase 1 (*gr1-1, gr1-2*) show a less negative cytosolic *E*_GSH_ than the wild type (WT) (Marty *et al*. 2009) with enhanced oxidation after osmotic stress (300 mM mannitol) (Bangash *et al*. 2019). In wildtype *A. thaliana*, oxidative shifts in cytosolic *E*_GSH_ have been observed under oxygen deprivation (Wagner *et al*. 2019), heat or heavy metal stress (Schwarzländer *et al*. 2009) and after herbicide treatment of chloroplasts (Ugalde *et al*. 2021).

While transgenic sensor lines in the model dicotyledon *A. thaliana* are in use since more than a decade, the progress in other genetically accessible model plants has only recently started to gain momentum (Kirschner et al. 2018; Müller-Schüssele et al. 2020; Hipsch et al. 2021; Giridhar et al. 2022). To start filling this gap in temperate cereal crops, we created transgenic barley lines expressing the genetically-encoded redox biosensor cytosolic Grx1-roGFP2 for *E*_GSH_. We aimed to investigate stress resilience in barley and to monitor *E*_GSH_ changes in different tissues *in vivo*. In this article, we characterise the obtained sensor lines and expose them to different stress treatments, revealing remarkable robustness of the cytosolic *E*_GSH_ in barley (cv. Golden Promise Fast).

## MATERIAL AND METHODS

### Plant materials and growth conditions

Experiments were conducted in “Golden Promise Fast”, the *Ppd-H1* introgression line of the transformable barley (*Hordeum vulgare* L.) cultivar Golden Promise (Gol *et al*. 2021; Amanda *et al*. 2022). After 3 days of stratification at 4°C on wet filter paper, seeds were transferred to germination paper (Anchor Paper Co, Saint Louis, USA). Barley was grown for up to 7 days in germination paper rolls with ½ strength Hoagland solution (Hoagland and Arnon, 1938). Stress treatments were carried out as described in Osthoff *et al*. (2019) with three days of growth in ½ strength Hoagland solution before transfer to nutrient solution with 150 mM NaCl and −0.8 MPa water potential (adjusted with PEG8000). Paper rolls were grown upright in a beaker in a climate cabinet under 16 h light (120 μmol photons m^−2^ s^−1^) at 22°C and 8 h dark at 18°C.

For propagation and seed production, 7-10-day-old barley seedlings were transferred to the green house and soil-grown with 15 h of light (1000-6000 lux) at 22°C and 9 h of dark at 18°C.

### Generation of barley Grx1-roGFP2 lines

A barley-compatible expression vector for Grx1-roGFP2 was constructed by restriction enzyme-based cloning in the binary vector p6i-2×35s-TE9, kindly provided by Jochen Kumlehn (Himmelbach *et al*. 2007), along with the pUBI-ABM (Watanabe *et al*. 2016), as source of the maize (*Zea mays*) ubiquitin promoter *ZmUbi1* and the *nos* terminator. A Grx1-roGFP2 fragment with BamHI and SalI restriction sites was ligated into the linearized UBIp-ABM (BamHI, Sall) with the ClonJet kit (ThermoFisher), to produce pZmUBI1pro-Grx1roGFP2-nosT. The ZmUBI1pro-Grx1roGFP2-nosT cassette was released by digestion with SfiI and ligated into binary vector p6i-d35STE9 previously cut with the same enzyme. The resulting expression plasmid was transformed in Golden Promise Fast as described in Amanda *et al*. (2022) and T_0_ hemizygous regenerants from 8 independent events were produced. We screened the T_1_ progenies from 5 out of the 8 independent events for roGFP2 fluorescence with a confocal laser scanning microscope and selected lines with 3:1 segregation of positive:negative fluorescence. T_2_ progenies of positive-fluorescent plants were further screened for 100% fluorescence expression to obtain homozygous transgenics. This procedure resulted in selecting two independent lines (5-3#39 and 2-1#1). We conducted all reported experiments in homozygous T_3_ progenies of these lines.

### Excitation scan of young barley leaves

Leaf discs were transferred into the wells of a 96-well plate containing 200 μL imaging buffer (10 mM MES, 5 mM KCI, 10 mM CaCl_2_, 10 mM MgCl_2_ pH 5.8). For reduction and oxidation of Grx1-roGFP2 the imaging buffer was supplemented with 1.5 M H_2_O_2_, 5 mM 2,2’-dipyridyl disulfide (DPS) or 10 mM dithiothreitol (DTT) to obtain total reduction and oxidation of the sensor. Leaf discs were vacuum infiltrated for 10 min and further incubated for 30 min. Fluorescence spectra were recorded with a plate reader (Clariostar®, BMG) with excitation wavelengths from 386-495 nm and emission set to 530/40 nm. Fluorescence was normalized over the fluorescence intensity at the isosbestic point (IP) of roGFP2 at 425 nm.

### Confocal laser scanning microscopy of Grx1-roGFP2

Confocal laser scanning microscopy of Grx1-roGFP2 in barley roots and leaves was conducted with a LSM780 (Axio Observer.Z1, Carl Zeiss, Oberkochen, Germany) using a 25x (Plan-Apochromat 25x/0.8) or 40x (C-Apochromat 40x/1.2W) objective by exciting roGFP2 at 405 nm (diode laser, 3%) and 488 nm (argon laser (1.5 %) and collecting roGFP2 fluorescence between 508 and 535 nm. Autofluorescence was detected at 430 to 470 nm after excitation at 405 nm. Ratio calculations and further image analysis was performed in a MATLAB-based ratio software (RRA) (Fricker 2016). Where indicated, regions of interest were set to nuclei as shown in **Suppl. Fig. 1**. Oxidation degree of roGFP2 (OxD) was calculated according to Schwarzländer *et al*. (2008).

### Glutathione detection in barley roots

After 96 h of stress treatment (see above), roots were incubated in 100 μM MCB (monochlorobimane) for exactly 30 min to label glutathione and subsequently transferred to a 50 μM propidium iodide (PI) solution for 5 min to label cell walls and prove cell viability (Meyer *et al*. 2001). After the labelling procedure, residual dyes were washed out in deionised water. Fluorescence was imaged after excitation at 405 nm at emission wavelengths of 449-613 nm (MCB) and after 543 nm excitation with emission wavelengths of 613-704 nm (PI). Confocal z-stacks were taken with identical settings with the confocal laser scanning microscope covering 270 μm of root tissue in 10 μm slices/image. Gain and laser settings were kept constant throughout the imaging process. Only roots showing no severe cell damage indicated by PI staining were further analysed in ImageJ as 8-bit images of maximum intensity projections of the z-stacks. The mean pixel intensity was extracted of the area of the root tip in each image and blotted as boxplot using GraphPad Prism 9.

### Glutathione reductase activity assay in barley root extracts

Flash-frozen and pulverized (TissueLyser (Qiagen) for 1.5 min, 30000 Hz) root material was dissolved in 100 μL of 100 mM K_2_HPO_4_/KH_2_PO_4_ at pH 7.5 supplemented with 0.5 mM EDTA and 0.1% of Plant Protease Inhibitor Cocktail P9599 (Sigma). Samples were vortexed and centrifuged at 20.000 x g 5 min at 4°C. The supernatant was transferred into a new tube and the centrifugation step was repeated. Protein concentration was determined in a Bradford assay (Bradford 1976). Extract containing 5 μg of protein was used in a total assay volume of 250 μL for the DTNB (5,5’-dithiobis(2-nitrobenzoic acid))-based GR activity assay (Smith *et al*. 1988; Marty *et al*. 2009), together with 1 mM EDTA, 750 μM DTNB, 1 mM GSSG in 100 mM K_2_HPO_4_/KH_2_PO_4_ at pH 7.5. TNB absorbance was followed at 412 nm using a plate reader (Clariostar®, BMG). After 5 min of measuring background activity, NADPH was added to a final concentration of 200 μM. Glutathione reductase activity was calculated via the increase of A_412nm_/min and the molar extinction coefficient of TNB.

## RESULTS

### Generation of stable barley lines expressing cytosolic Grx1-roGFP2

We generated barley (cv. Golden Promise Fast) lines constitutively expressing Grx1-roGFP2 under the control of the maize ubiquitin promotor (*ZmUbi*), including the 5’UTR intron. Two independent lines originating from shoots from different calli were chosen based on consistent GFP fluorescence (5-3 #39 and 2-1 #1) and brought to homozygosity. In the chosen homozygous lines, we observed Grx1-roGFP2 signal, independent of tissue type or developmental stage (**Suppl. Fig. 1**). No phenotypic difference in growth was observed between barley expressing the sensor and the Golden Promise Fast background.

The epidermal layer of barley leaves showed the typical sensor fluorescence signal in the nucleus and cytosol **(Fig. 1A)**. High autofluorescence signal was detected after excitation at 405 nm in guard cells and in vacuoles of mesophyll cells, as well as in the wall of root cells.

**Figure 1:**
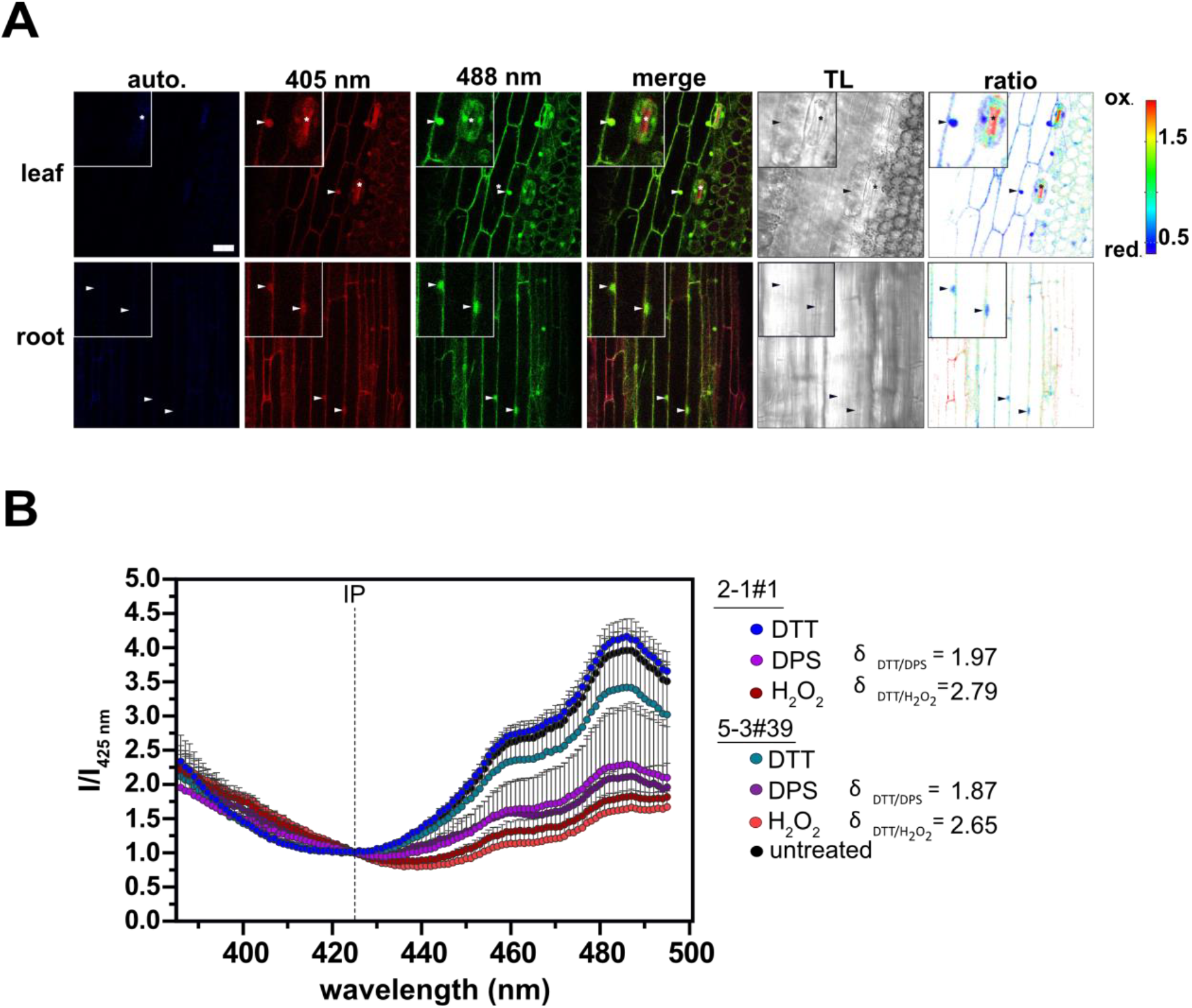
Grx1-roGFP2 expression in barley leaf and root tissue. **A** Confocal microscopy images of roGFP2 (405 nm and 488 nm excitation; 508-535 nm emission) and autofluorescence (auto, 405 nm excitation, 430-470 nm emission). The merge image displays the overlay of 405 and 488 intensities. TL displays the transmitted light image. Arrowheads indicate nucleus- and cytosol-localized sensor signal, and the asterisks show regions of high autofluorescence. Scale bar = 40 μm. **B** Excitation scan (386-495 nm; emission: 530-40 nm) of barley leaf discs treated with 10 mM DTT, 5 mM DPS or 1.5 M H_2_O_2_. Fluorescence was normalized over the fluorescence intensity at the isosbestic point (IP) of roGFP2 at 425 nm. Significance between genotypes and treatments of the 405 and 488 nm peak intensity was tested with a Two-way ANOVA and Tukey’s post hoc test with α<0.05. No significant differences were observed between the lines within one treatment while peak intensities of 488 nm were significantly different in reducing (DTT) and oxidizing (H_2_O_2_ and DPS) treatments.

### Cytosolic Grx1-roGFP2 reacts to oxidising and reducing agents in barley shoots and roots

To assess if the sensor is responsive to exogenously induced reduction or oxidation, leaf discs of 7-day-old barley plants were exposed to oxidising and reducing agents. We observed a roGFP2-typical *in planta* excitation spectrum after excitation of the leaf discs at 386 to 495 nm (**Fig. 1B**). Reduction of the sensor with DTT lead to a higher excitation peak while oxidation with H_2_O_2_ or DPS lead to a lower excitation peak in the spectral range above the isosbestic point (IP) at 425 nm. Below the IP, the influence of sensor oxidation or reduction on *in vivo* excitation spectra was only low, resulting in a minor contribution to 405/488 ratio changes. The *in vivo* spectroscopic dynamic range (δ) of Grx1-roGFP2 was calculated to be around 2- to −3-fold (405/488) (×2.79 2-1 #1; ×2.65 5-3 #39). Untreated barley leaves showed a similar spectral signal as the DTT-treated leaves, suggesting a largely reduced sensor under physiological conditions. Furthermore, the sensor did not behave significantly different between the two independent lines.

We further determined the *in vivo* dynamic range of Grx1-roGFP2 in barley plants based on confocal microscopy. As the nuclear roGFP2 signal in root and leaf tissue was not affected by interfering autofluorescence signals we used nuclear regions of interest during image analysis to minimize cell wall and vacuole fluorescence interference (**Fig. 2, Suppl. Figs. 2 and 3**). We calibrated Grx1-roGFP2 in root and leaf tissue of young barley seedlings and observed tissue-dependent differences (**Suppl. Fig. 2**). While the calibration of line 5-3 #39 and line 2-1 #1 in leaves resulted in a dynamic range of δ = 1.3-1.5, calibration of the root tissue resulted in a dynamic range of δ = 2.3-2.7 depending on the used oxidizing agent. Furthermore, the physiological state of Grx1-roGFP2 in roots revealed a higher 405/488 nm ratio of 0.93 ± 0.50 (2-1 #1) and 1.05 ± 0.61 (5-3 #39) compared to leaves with a ratio of 0.63 ± 0.27 (2-1 #1) and 0.60 ± 0.18 (5-3 #39), suggesting a slightly more oxidized sensor in roots than in leaves.

**Figure 2:**
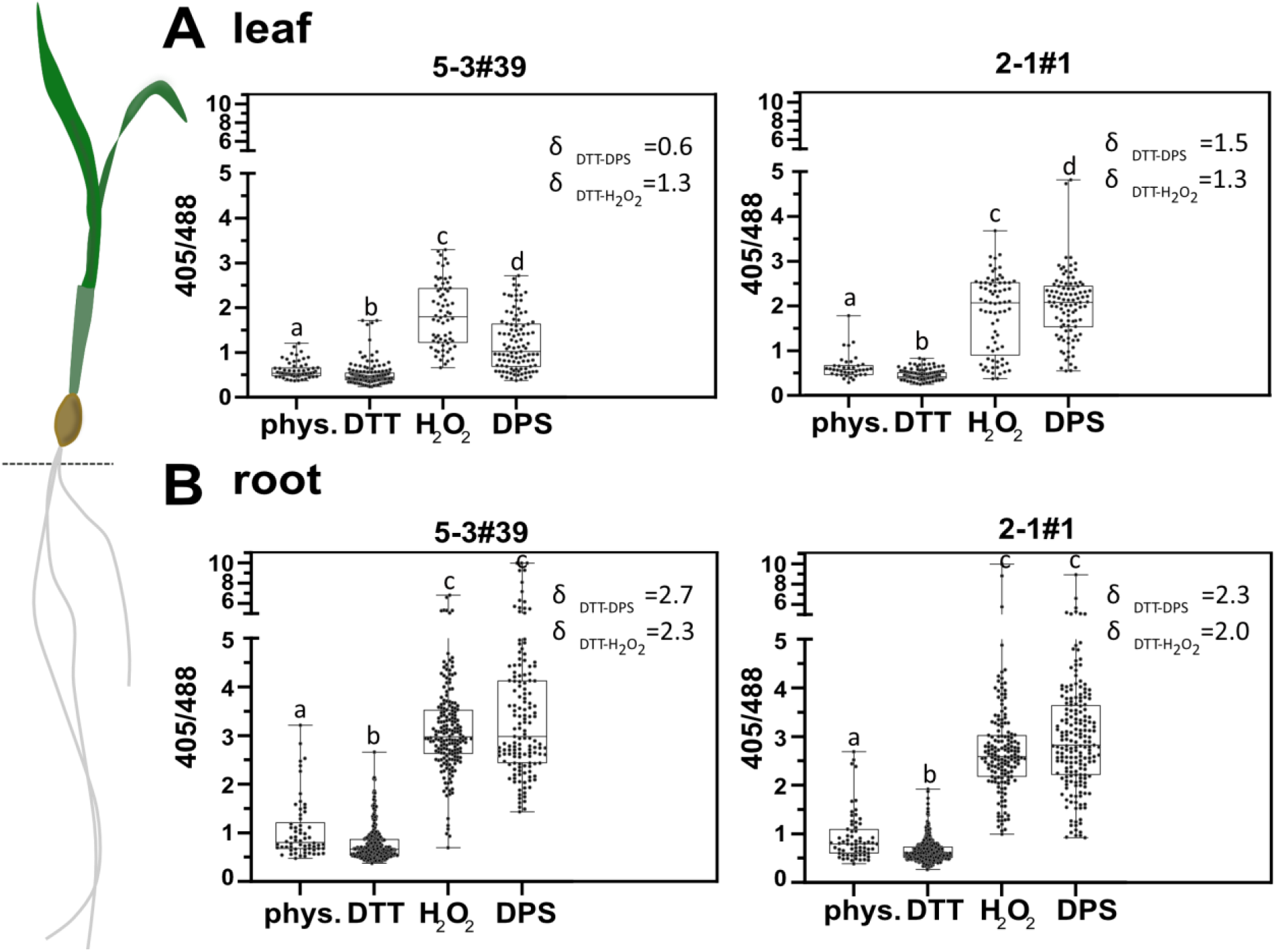
Grx1-roGFP2 sensor calibration in barley leaf and root tissue. Box plots of Grx1-roGFP2 signal ratios (405/408) calculated from fluorescence intensities in nuclei of leaf (**A**) and root (**B**) tissue treated with 10 mM DTT, 5 mM DPS or 1.5 M H_2_O_2_ and imaged via confocal microscopy. A one-way ANOVA and Tukey’s multiple comparison test was conducted on log transformed ratio values. Significantly different treatments (p < 0,0001) are indicated by different letters. Boxes display 25 to 75 percentiles with whiskers showing min to max values. The middle line indicates the median and dots represent the data of every nucleus obtained from 2 to 5 seedlings per treatment.

### Cytosolic *E*_GSH_ reacts only slowly and mildly to severe growth-impairing osmotic and salt stress

A recent transcriptomic study (Osthoff *et al*. 2019) showed that barley seedlings treated with a combination of salt and osmotic stress in a paper roll system upregulate transcript encoding the cytosolic isoform of glutathione reductase (GR1; HORVU6Hr1G089780) after 24 h of stress exposure. As the expression of the most important enzyme for maintaining a highly reduced cytosolic glutathione redox status was affected under these conditions and the used paper roll system allows for soil-free imaging of barley plants, we chose the same experimental conditions for barley expressing Grx1-roGFP2 to assess potential effects of this stress treatment on the cytosolic *E*_GSH_.

As reported by Osthoff *et al*. (2019), barley seedlings treated with a combined stress of 150 mM NaCl and a water potential of −0.8 MPa (adjusted using PEG8000) showed a decrease in root growth (**Fig. 3**). While the mean root growth of barley grown for 7 days in ½ strength Hoagland solution showed root lengths between 57.47 ± 22.05 mm (mean of 5-3 #39 and 2-1 #1, control), root growth decreased significantly after a combined salt and osmotic stress for 96 h to 34.7 ± 14.81 mm (mean of 5-3 #39 and 2-1 #1, 150 mM NaCl, −0.8 MPa PEG8000).

**Figure 3:**
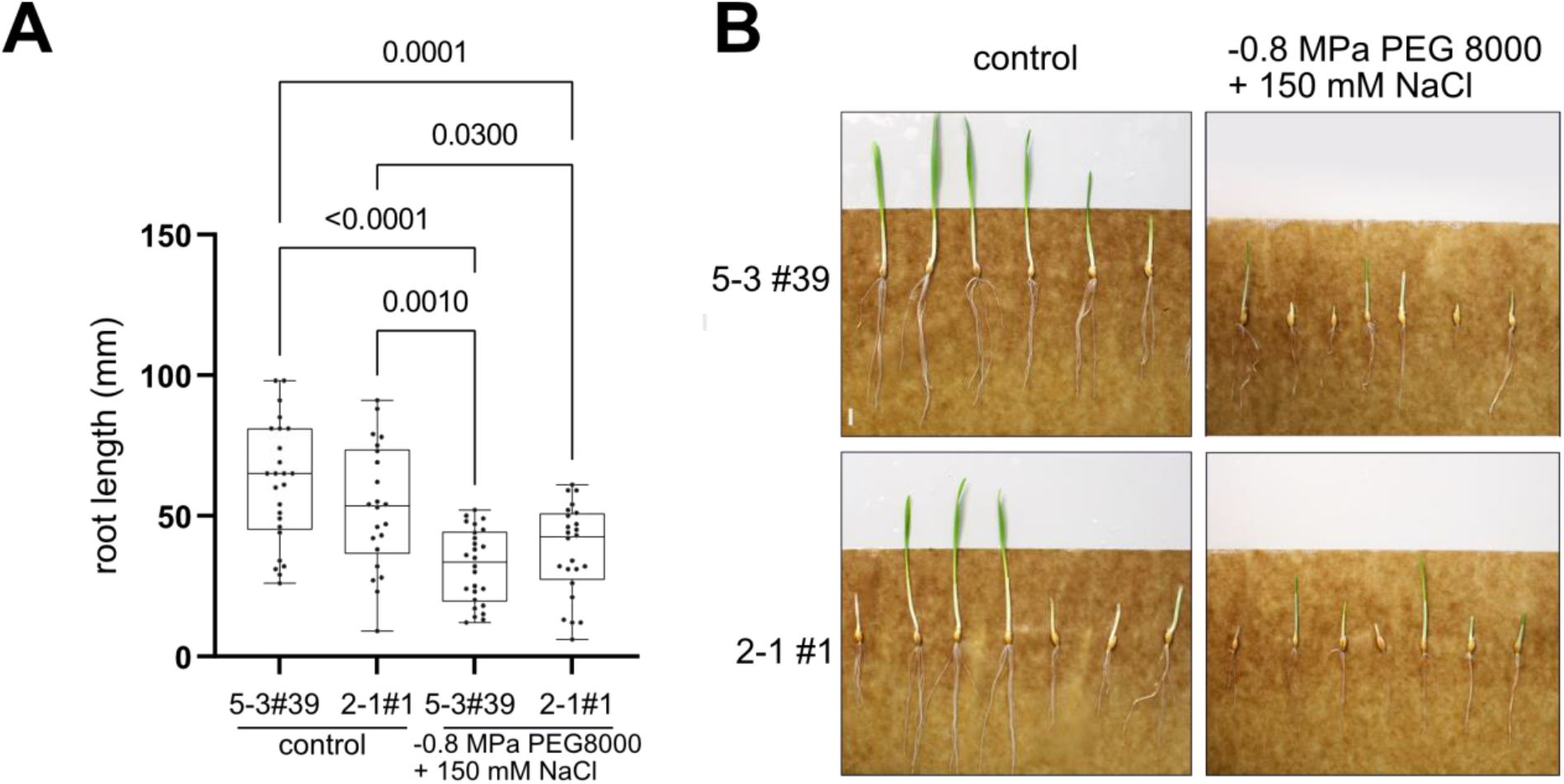
Effect of combined osmotic and salt stress on barley seedling root growth. **A** Root length of 7-day old barley plants after 96 h of combined osmotic and salt stress. Boxes display 25 to 75 percentiles with whiskers showing min to max values. The median is indicated by the horizontal line. Dots show individual root lengths measured of 21-26 individual seedlings. **B** Example images of Grx1-roGFP2 sensor lines under control and stress conditions, 96 h after beginning of the stress.

Next, we followed the redox state of the Grx1-roGFP2 sensor at 24 h, 48 h and 96 h after the start of the stress (Osthoff *et al*. 2019). Grx1-roGFP2 redox state was read out via confocal laser scanning microscopy and quantified via ratio image analysis of nuclei as regions of interest (ROIs). We found that cytosolic *E*_GSH_ is maintained during the first 48 After 96 ha significant but small oxidative shift in the 405/488 nm ratio between stressed (1.05) and non-stressed (0.86) plants (**Fig. 4, Suppl. Figs. 4, 5, 6**). The measured difference in 405/488 nm ratio corresponds to only a 3% shift in degree of sensor oxidation.

**Figure 4:**
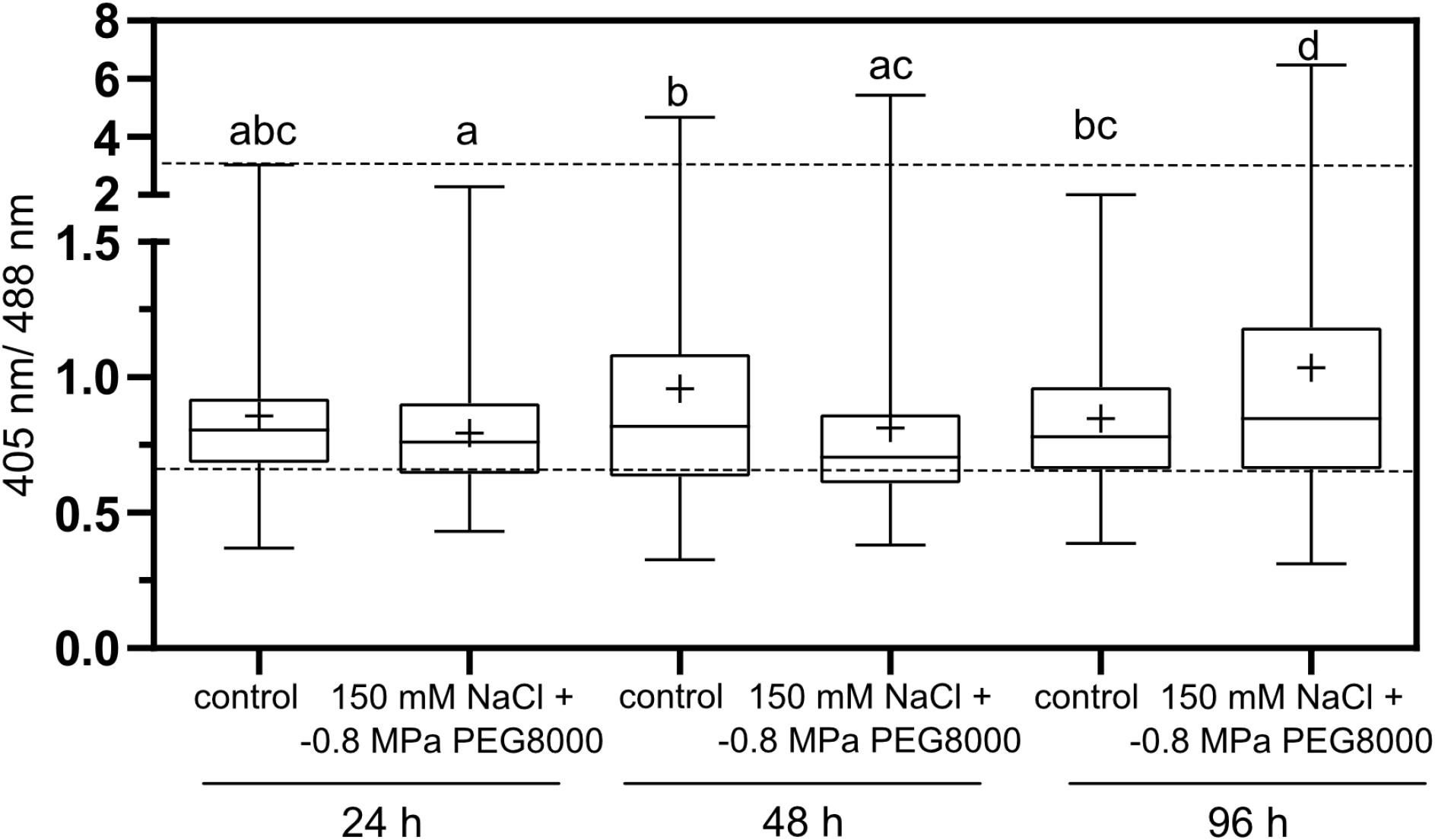
roGFP2 ration in roots of barley seedlings after 24, 48 or 96 h of combined osmotic and salt stress. Box plots of roGFP2 nuclear signal ratio (405/488). Boxes display 25 to 75 percentiles with whiskers showing min to max values. The median is displayed as horizontal line and the mean is indicated with the ‘+’ sign. Dotted lines indicate 100 % oxidation and reduction of the sensor (calibration with line 2-1 #1 root values, see **Fig. 2**). Two-way ANOVA was conducted on log-transformed values with α = 0.05, different letters indicate significantly different categories. Between 170-256 nuclei were analysed from 3-5 seedlings per line and treatment.

### Investigating the molecular basis for *E*_GSH_ robustness under stress

According to the Nernst equation, *E*_GSH_ is dependent on both the total concentration of GSH and the amount of glutathione disulfide (GSSG). Thus, a less negative local *E*_GSH_ as measured via roGFP2 redox state could be caused by either increased GSSG levels or decreased total glutathione content in the cytosol. Glutathione reductase recycles GSSG back to GSH, using NADPH as electron donor. As the K_m_ of this important enzyme is in the nanomolar range, [GSSG] would only rise if enzymatic capacity of GR is not sufficient to reduce GSSG immediately or if there is a shortage of NADPH. Transcriptomic analyses of Osthoff *et al*. (2019) identified the transcripts of both barley GR isoforms as differentially upregulated after 24 h of combined salt and osmotic stress (GR1: HORVU6Hr1G089780 log_2_FC 1.3; GR2: HORVU4Hr1G073930 log_2_FC 1.5). In contrast, transcripts encoding barley orthologs of the two proteins involved in GSH biosynthesis (GSH1 HORVU1Hr1G015590 and GSH2 HORVU5Hr1G027100) were not differentially expressed under those conditions (Osthoff *et al*. 2019).

Thus, we used an enzymatic assay for GR activity in plant extracts to test if total GR activity is increased in response to the same stress treatment. We did not find significantly reduced or increased total GR activity in the 96 h stress treatment samples (**Fig. 5 A**). The GR activity of the 2-1 #1 line under control and stress conditions remained constant with values of 105.5 ± 36.4 and 105.9 ± 39.8 nmol TNB/min/mg protein, similar to the 5-3 #39 line with a slightly lower measured GR activity of 98.9 ± 46.7 and 92.3 ± 43 nmol TNB/min/mg protein. To compensate for possible differences in the individual measurements, we normalized the measured activities after stress treatment with those of the corresponding control treatment (**Fig. 5 B**). The resulting normalized ratio values of 1.1 ± 0.5 (2-1 #1) and 1.0 ± 0.6 (5-3 #39) did not show significant differences in response to the stress treatment and maintained a high variance.

**Figure 5:**
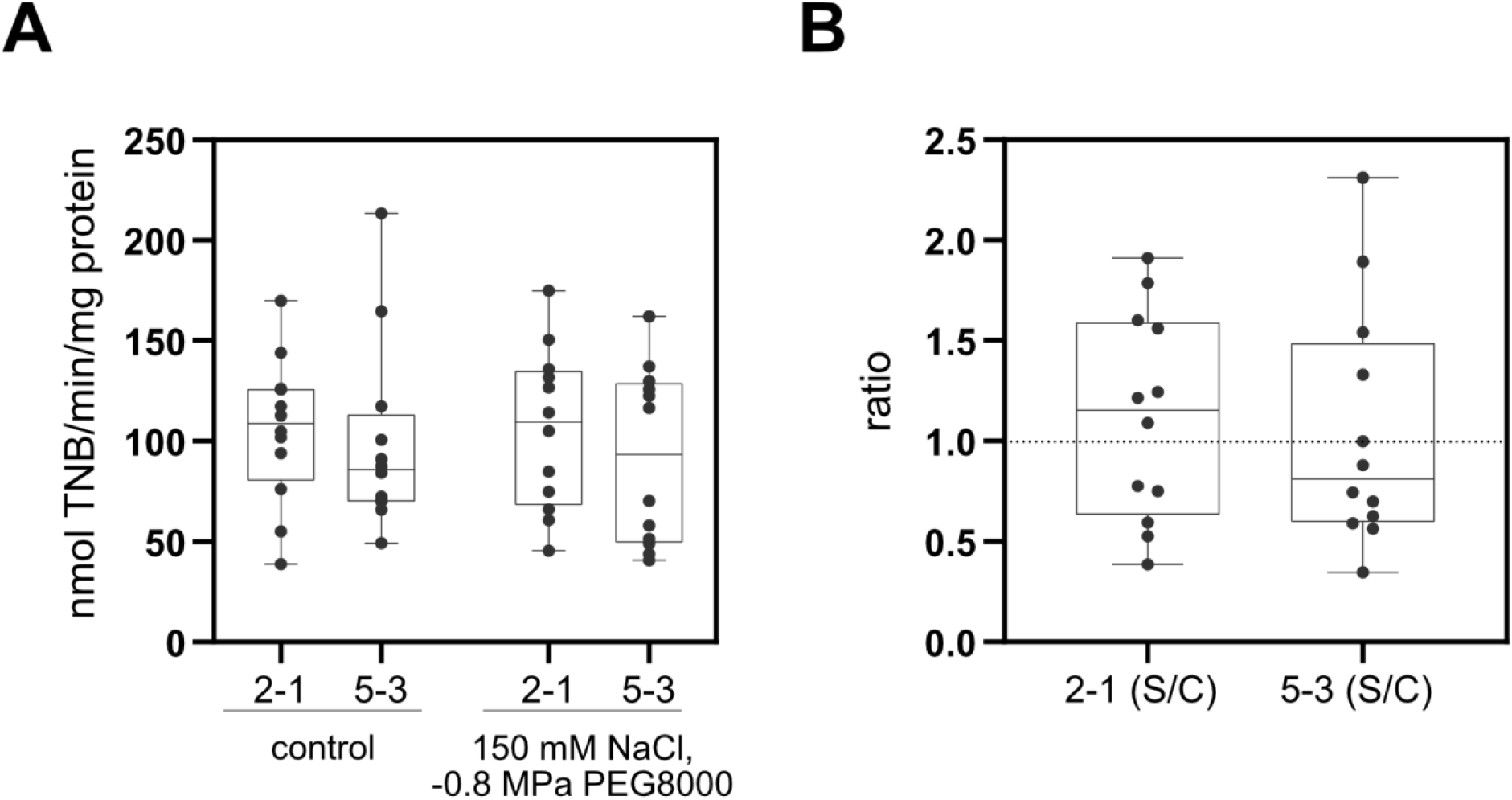
DTNB – based glutathione reductase assay in barley roots after osmotic and salt stress. **A** Absolute GR activity under control and 96 h of stress conditions displayed as box plots with boxes displaying 25 to 75 percentiles and whiskers showing min to max. Median is shown as horizontal line. Individual GR activities of 12 biological replicates for each line and condition are shown as dots. **B** GR activity in stressed samples was divided by the corresponding control activity. A Two-way ANOVA (A) and T-test (B) was conducted on data sets A and B with α = 0.05 and no significant differences were observed.

We additionally tested if the stress treatment modifies the total glutathione content after 96 h (**Fig. 6**). emitting fluorescence at 449-613 nm after excitation at 405 nm; Haugland *et al*. 1996). MCB-stained roots of stressed (150 mM NaCl, −0.8 MPa PEG8000, 96 h) and unstressed seedlings were imaged with a confocal laser scanning microscope., excluding a large increase in glutathione content after stress treatment

**Figure 6:**
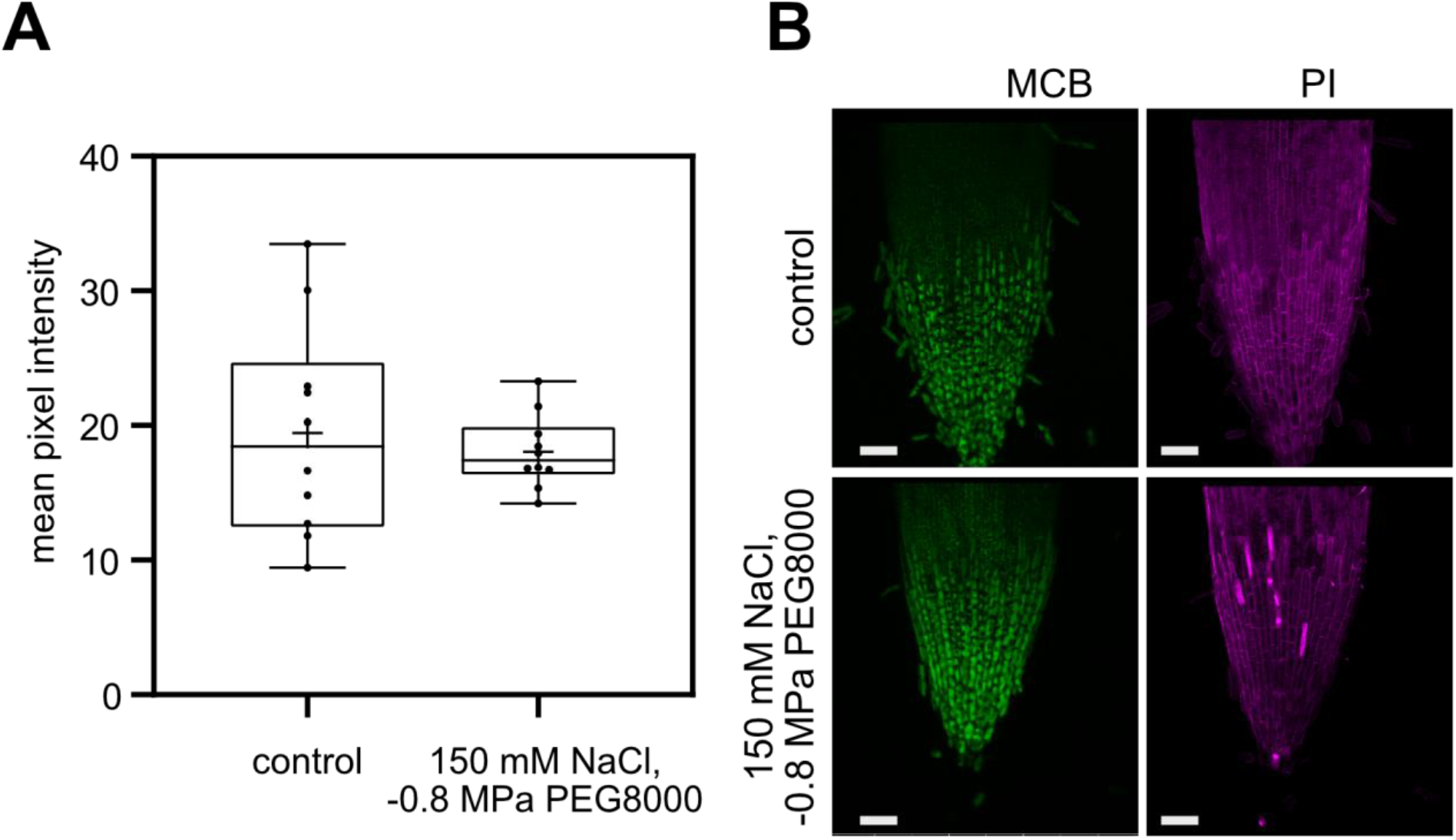
*In situ* detection of total glutathione and cell damage in stressed barley root tips (96 h). **A** Mean pixel intensities of root tips stained with MCB were extracted with ImageJ (8-bit, 0-255 display range). Boxes display 25 to 75 percentiles with whiskers showing min to max values, the median is displayed as constant line. The mean of all data points is shown as ‘+’ (n = 10). No significant differences were detected (T-test). **B** Maximum intensity projection of barley root tips stained with monochlorobimane (MCB) (excitation: 405 nm, emission: 449-613 nm) and propidium iodide (PI, excitation: 543 nm, emission: 613-704 nm). Scale bar = 100 μm.

## DISCUSSION

In this study we generated and characterised two independent transgenic barley (*Hordeum vulgare*, cv. Golden Promise Fast) lines expressing the genetically encoded biosensor Grx1-roGFP2 under the control of the *Zea mays* Ubiquitin promoter including the 5’ UTR intron (Christensen & Quail 1996). As expected, the biosensor was constitutively expressed and localised to the cytosol and nucleus. The fluorescence intensity was sufficiently high for ratiometric sensor read-out in all examined tissues. As we observed high autofluorescence after excitation with UV light (405 nm) in the apoplast and sometimes the vacuole and cell walls, we preferably calculated 405/488 nm ratios from regions of interest set to nuclei. Using this approach, we determined that the *in vivo* dynamic range of cytosolic Grx1-roGFP is c. 1.3 - 1.5 in leaves and c. 2.3 - 2.7 in roots. The previously reported *in vitro* dynamic range of roGFP2 was c. 8.2 - 9.2 (405/488 nm) (Schwarzländer *et al*. 2008; Aller *et al*. 2013) while *in vivo* samples often displayed decreased dynamic ranges of c. 5 in confocal microscopy or c. 3 in plate reader-based sensor read-out (Ugalde *et al*. 2021, 2022). This difference in dynamic range between *in vitro* and *in vivo* sensor read-out can be explained by interference of plant compounds that overlay the changes in sensor absorption in the UV range (below the isosbestic point at c. 425 nm). This is also the case for barley, where the change in roGFP2 excitation in the UV range between fully reduced and fully oxidized plant samples is very small (**Fig. 1B**). Thus, the 405/488 sensor ratio displays a lower dynamic range *in vivo* than *in vitro*. The observed differences in calibration between leaf samples and root samples (**Fig. 2**) could be attributed to insufficient penetration of the oxidants H_2_O_2_ and DPS into barley leaves. However, within each tissue sensor oxidation caused by 1.5 M H_2_O_2_ or 5 mM DPS were not significantly different. Usually, lower concentrations of H_2_O_2_ (10-100 mM) are used for *in vivo* roGFP2 sensor calibration but were not effective in barley (this study) and potato (Hipsch *et al*. 2021). Generally, it is very important to identify the optimal conditions for sensor calibration that allow both determination of the 405/488 nm ratio values for 0% and 100% roGFP oxidation (OxD), especially when mV values for the glutathione redox potential are to be calculated (reviewed in Müller-Schüssele *et al*. 2021, Schwarzländer, *et al*. 2021). Using the sensor calibration in roots and the untreated (physiological) root samples, we calculated that the cytosolic *E*_GSH_ of the barley cytosol is c. −308 to −320 mV assuming a pH of 7.2 (**Suppl. Fig. 2**). This is close to the cytosolic *E*_GSH_ in *A. thaliana*, which was measured as c. −320 to −310 mV (Meyer *et al*. 2007; Schwarzländer *et al*. 2008; Aller *et al*. 2013). This *E*_GSH_ in the barley cytosol results in mostly reduced Grx1-roGFP2, supporting the suitability of Grx1-roGFP2 to sense oxidative changes in this compartment.

We also observed a small, but statistically significant difference in the 405/488 nm sensor ratio between barley roots and leaves (**Fig. 2, Suppl. Fig. 2**), which suggests a slightly more reduced physiological steady state of Grx1-roGFP2 in leaves than in roots, corresponding to a 5-10 mV difference. However, we cannot rule out that read-out of roGFP2 excitation ratios differs slightly between these organs due to tissue-specific differences in wavelength penetration or autofluorescence.

Many molecules in plants emit fluorescence after excitation in the UV range such as chlorophyll, lignin or alkaloids (Donaldson, 2020). We detected high levels of autofluorescence in barley leaves and roots excited with 405 nm. Since cells may contain multiple autofluorescent compounds, we cannot narrow down the exact source of autofluorescence in these tissues after excitation with 405 nm. Barley tissues are rich in phenolic compounds such as the phenylpropanoid ferulic acid (4-hydroxy-3-methoxy-cinnamicacid) with concentrations ranging from 359-624 μg/g dry weight (Hernanz et al., 2001; Bonoli et al., 2004).

By applying stress to the generated reporter lines, we observed shorter seminal root lengths after 96 h (4 days) of combined salt and osmotic stress comparable to Osthoff *et al*. (2019). We therefore used this time point to asses shifts in the cytosolic glutathione redox potential. We first monitored the roGFP2 redox state 24 h, 48 h and 96 h after combined stress treatment and where able to detect significant but small changes in the roGFP2 oxidation degree after 96 h. The measured shift in roGFP2 405/488 nm ratio corresponds to a 3% shift in degree of oxidation, implying a + 2.1 mV shift from −314 mV (control 96 h) to −311.9 mV (150 mM NaCl, −0.8 MPa PEG8000) (pH 7.2, calibration values of 2-1 #1 root).

Water deficit and salt stress caused upregulation of transcripts related to the response to an oxidative challenge (Osthoff *et al*. 2019). Several enzymes involved in the response to oxidative stress draw electrons from the glutathione pool, generating local GSSG. The highest flux of electrons can arguably be expected from DHAR, which re-reduces dehydroascorbate to ascorbate in the ascorbate/glutathione cycle that scavenges H_2_O_2_ (Foyer & Noctor 2011). As we detected only a slight oxidative response of Grx1-roGFP2 after 96 h of combined water deficit and salt stress, the cytosolic *E*_GSH_ in barley seems highly robust against oxidative changes. Possible reasons could be that (1) GSH does not serve as electron donor to mitigate these stress conditions or (2) GSH serves as electron donor and generates GSSG but compensatory mechanisms keep *E*_GSH_ constant. According to the Nernst equation, *E*_GSH_ could be kept constant via an increase in total GSH or removal of GSSG. To test these scenarios, we assessed total glutathione content and total glutathione reductase activity, but we found no changes in these parameters under the tested stress conditions. However, it should be noted that our imaging-based approach is limited regarding the investigated tissue type and that the enzyme activity-based approach can only assess total glutathione reductase activity (i.e. GR1 and GR2). Thus, we cannot completely rule out an increase in activity of cytosolic GR1 under osmotic and salt stress. Another, untested possibility is an efficient export of GSSG from the cytosol, e.g. to the vacuole (Morgan *et al*. 2013; Marty *et al*. 2019).

We created our reporter lines in the transformable background Golden Promise Fast, which has shown resilience under drought stress (Gol *et al*. 2021). Our lines in this background showed reductions in root growth under combined salt and osmotic stress, similarly to the German spring barley cultivar Scarlett (Osthoff *et al*. 2019), indicating that Golden Promise Fast does respond to these abiotic stresses. However, it is formally possible that the robust redox state displayed by Golden Promise Fast under our combined stress treatments are a particularity of this genetic background. Thus, it may be of interest to introgress the reporter in other cultivars and study its response to similar stresses.

In conclusion, we generated barley reporter lines that allow to monitor the cytosolic *E*_GSH_ via Grx1-roGFP2 oxidation state and that can be used to further investigate stress tolerance in barley. To further dissect if the high robustness of cytosolic *E*_GSH_ correlates with high stress resilience further investigations will be required, such as genetic analyses of different redox enzyme mutants.

In general, the glutathione-dependent redox system is viewed as a house-keeping redox system and *E*_GSH_ as a relatively constant parameter in the plant cytosol. Given the robustness of *E*_GSH_, it may be tempting to propose that this parameter could be a strong predictor of irreversible damage and imminent cell death. In this regard, *E*_GSH_ reporter lines would be interesting resources to screen for higher stress resilience.

## Author Contributions

FB, IA, AJM and SJMS designed the research. FB, AK and HT performed experiments and analysed data. SJMS, MS, AJM, FH and IA supervised the research and provided resources.

FB and SJMS wrote the manuscript with contributions from all authors. All authors approved the manuscript before submission.

## Acknowledgements

We thank Alexa Brox and Maria Homagk (Chemical Signalling, INRES, University of Bonn) for assistance in cloning, barley cultivation and stress treatments, and Edelgard Wendeler (MPIPZ) for her technical support for the generation of the transgenic lines.

This work was supported by the DFG-funded Research Training Group GRK2064: “Water use efficiency and drought stress responses: From Arabidopsis to Barley” (AJM,. MS, FAB and SJM), and the Max Planck Society (IFA).

## Literature

Aller I., Rouhier N., Meyer A.J. (2013) Development of roGFP2-derived redox probes for measurement of the glutathione redox potential in the cytosol of severely glutathione-deficient *rml1* seedlings. Frontiers in Plant Science 4:506.

Amanda D., Frey F.P., Neumann U., Przybyl M., Šimura J., Zhang Y., Chen Z., Gallavotti A., Fernie A.R., Ljung K., Acosta I.F. (2022) Auxin boosts energy generation pathways to fuel pollen maturation in barley. Current Biology 32:1798–1811.e8.

Bangash S.A.K., Müller-Schüssele S.J., Solbach D., Jansen M., Fiorani F., Schwarzländer M., Kopriva S., Meyer A.J. (2019) Low-glutathione mutants are impaired in growth but do not show an increased sensitivity to moderate water deficit. PloS One 14:e0220589.

Bradford M.M. (1976) A rapid and sensitive method for the quantitation of microgram quantities of protein utilizing the principle of protein-dye binding. Analytical Biochemistry 72:248–254.

Christensen A.H., Quail P.H. (1996) Ubiquitin promoter-based vectors for high-level expression of selectable and/or screenable marker genes in monocotyledonous plants. Transgenic Research 5:213–218.

Dawson I.K., Russell J., Powell W., Steffenson B., Thomas W.T.B., Waugh R. (2015) Barley: a translational model for adaptation to climate change. New Phytologist 206:913–931.

Foyer C.H., Noctor G. (2011) Ascorbate and glutathione: the heart of the redox hub. Plant Physiology 155:2–18.

Fricker M.D. (2016) Quantitative redox imaging software. Antioxidants & Redox Signaling 24:752–762.

Giridhar M., Meier B., Imani J., Kogel K.-H., Peiter E., Vothknecht U.C., Chigri F. (2022) Comparative analysis of stress-induced calcium signals in the crop species barley and the model plant *Arabidopsis thaliana*. BMC Plant Biology 22:447.

Gol L., Haraldsson E.B., von Korff M. (2021) Ppd-H1 integrates drought stress signals to control spike development and flowering time in barley (M. Jones, Ed.). Journal of Experimental Botany 72:122–136.

Gutscher M., Pauleau A.-L., Marty L., Brach T., Wabnitz G.H., Samstag Y., Meyer A.J., Dick T.P. (2008) Real-time imaging of the intracellular glutathione redox potential. Nature Methods 5:553–559.

Haugland R.P., Spence M.T.Z., Johnson I.D. (1996) Handbook of fluorescent probes and research chemicals, 6th ed. Molecular Probes, Eugene, OR, USA (4849 Pitchford Ave., Eugene 97402).

Hoagland D.R, Arnon D.I. The water-culture method for growing plants without soil. College of Agriculture, University of California, 1938

Himmelbach A., Zierold U., Hensel G., Riechen J., Douchkov D., Schweizer P., Kumlehn J. (2007) A set of modular binary vectors for transformation of cereals. Plant Physiology 145:1192–1200.

Hipsch M., Lampl N., Zelinger E., Barda O., Waiger D., Rosenwasser S. (2021) Sensing stress responses in potato with whole-plant redox imaging. Plant Physiology 187:618–631.

Imani J., Li L., Schäfer P., Kogel K.-H. (2011) STARTS - A stable root transformation system for rapid functional analyses of proteins of the monocot model plant barley: Stable barley root transformation system. The Plant Journal 67:726–735.

Kirschner G.K., Stahl Y., Imani J., von Korff M., Simon R. (2018) Fluorescent reporter lines for auxin and cytokinin signalling in barley (*Hordeum vulgare*) (X. Li, Ed.). PLOS ONE 13:e0196086.

Kreszies T., Shellakkutti N., Osthoff A., Yu P., Baldauf J.A., Zeisler-Diehl V.V., Ranathunge K., Hochholdinger F., Schreiber L. (2019) Osmotic stress enhances suberization of apoplastic barriers in barley seminal roots: analysis of chemical, transcriptomic and physiological responses. New Phytologist 221:180–194.

Marty L., Bausewein D., Müller C., Bangash S.A.K., Moseler A., Schwarzländer M., Müller-Schüssele S.J., Zechmann B., Riondet C., Balk J., Wirtz M., Hell R., Reichheld J.-P., Meyer A.J. (2019) Arabidopsis glutathione reductase 2 is indispensable in plastids, while mitochondrial glutathione is safeguarded by additional reduction and transport systems. The New Phytologist 224:1569–1584.

Marty L., Siala W., Schwarzlander M., Fricker M.D., Wirtz M., Sweetlove L.J., Meyer Y., Meyer A.J., Reichheld J.-P., Hell R. (2009) The NADPH-dependent thioredoxin system constitutes a functional backup for cytosolic glutathione reductase in Arabidopsis. Proceedings of the National Academy of Sciences 106:9109–9114.

Mascher M., Gundlach H., Himmelbach A., Beier S., Twardziok S.O., Wicker T., Radchuk V., Dockter C., Hedley P.E., Russell J., Bayer M., Ramsay L., Liu H., Haberer G., Zhang X.-Q., Zhang Q., Barrero R.A., Li L., Taudien S., Groth M., Felder M., Hastie A., Šimková H., Staňková H., Vrána J., Chan S., Muñoz-Amatriaín M., Ounit R., Wanamaker S., Bolser D., Colmsee C., Schmutzer T., Aliyeva-Schnorr L., Grasso S., Tanskanen J., Chailyan A., Sampath D., Heavens D., Clissold L., Cao S., Chapman B., Dai F., Han Y., Li H., Li X., Lin C., McCooke J.K., Tan C., Wang P., Wang S., Yin S., Zhou G., Poland J.A., Bellgard M.I., Borisjuk L., Houben A., Doležel J., Ayling S., Lonardi S., Kersey P., Langridge P., Muehlbauer G.J., Clark M.D., Caccamo M., Schulman A.H., Mayer K.F.X., Platzer M., Close T.J., Scholz U., Hansson M., Zhang G., Braumann I., Spannagl M., Li C., Waugh R., Stein N. (2017) A chromosome conformation capture ordered sequence of the barley genome. Nature 544:427–433.

Meyer A.J., Brach T., Marty L., Kreye S., Rouhier N., Jacquot J.-P., Hell R. (2007) Redox-sensitive GFP in *Arabidopsis thaliana* is a quantitative biosensor for the redox potential of the cellular glutathione redox buffer. The Plant Journal 52:973–986.

Meyer A.J., Dreyer A., Ugalde J.M., Feitosa-Araujo E., Dietz K.-J., Schwarzländer M. (2021) Shifting paradigms and novel players in Cys-based redox regulation and ROS signaling in plants - and where to go next. Biological Chemistry 402:399–423.

Meyer A.J., May M.J., Fricker M. (2001) Quantitative *in vivo* measurement of glutathione in Arabidopsis cells: *In vivo* measurement of glutathione. The Plant Journal 27:67–78.

Morgan B., Ezeriņa D., Amoako T.N.E., Riemer J., Seedorf M., Dick T.P. (2013) Multiple glutathione disulfide removal pathways mediate cytosolic redox homeostasis. Nature Chemical Biology 9:119–125.

Müller-Schüssele S.J., Bohle F., Rossi J., Trost P., Meyer A.J., Zaffagnini M. (2021) Plasticity in plastid redox networks: evolution of glutathione-dependent redox cascades and glutathionylation sites. BMC Plant Biology 21:322.

Müller-Schüssele S.J., Schwarzländer M., Meyer A.J. (2021) Live monitoring of plant redox and energy physiology with genetically encoded biosensors. Plant Physiology 186:93–109.

Müller-Schüssele S.J., Wang R., Gütle D.D., Romer J., Rodriguez-Franco M., Scholz M., Buchert F., Lüth V.M., Kopriva S., Dörmann P., Schwarzländer M., Reski R., Hippler M., Meyer A.J. (2020) Chloroplasts require glutathione reductase to balance reactive oxygen species and maintain efficient photosynthesis. The Plant Journal 103:1140–1154.

Munns R., James R.A., Läuchli A. (2006) Approaches to increasing the salt tolerance of wheat and other cereals. Journal of Experimental Botany 57:1025–1043.

Muzammil S., Shrestha A., Dadshani S., Pillen K., Siddique S., Léon J., Naz A.A. (2018) An ancestral allele of *pyrroline-5-carboxylate synthase1* promotes proline accumulation and drought adaptation in cultivated barley. Plant Physiology 178:771–782.

Newton A.C., Flavell A.J., George T.S., Leat P., Mullholland B., Ramsay L., Revoredo-Giha C., Russell J., Steffenson B.J., Swanston J.S., Thomas W.T.B., Waugh R., White P.J., Bingham I.J. (2011) Crops that feed the world 4. Barley: a resilient crop? Strengths and weaknesses in the context of food security. Food Security 3:141–178.

Osthoff A., Donà dalle Rose P., Baldauf J.A., Piepho H.-P., Hochholdinger F. (2019) Transcriptomic reprogramming of barley seminal roots by combined water deficit and salt stress. BMC Genomics 20:325.

Rey P., Tarrago L. (2018) Physiological roles of plant methionine sulfoxide reductases in redox homeostasis and signaling. Antioxidants 7:114.

Schwarzländer M., Dick T.P., Meyer A.J., Morgan B. (2016) Dissecting redox biology using fluorescent protein sensors. Antioxidants & Redox Signaling 24:680–712.

Schwarzländer M., Fricker M.D., MüLler C., Marty L., Brach T., Novak J., Sweetlove L.J., Hell R., Meyer A.J. (2008) Confocal imaging of glutathione redox potential in living plant cells. Journal of Microscopy 231:299–316.

Schwarzländer M., Fricker M.D., Sweetlove L.J. (2009) Monitoring the *in vivo* redox state of plant mitochondria: Effect of respiratory inhibitors, abiotic stress and assessment of recovery from oxidative challenge. Biochimica et Biophysica Acta (BBA) - Bioenergetics 1787:468–475.

Smith I.K., Vierheller T.L., Thorne C.A. (1988) Assay of glutathione reductase in crude tissue homogenates using 5,5’-dithiobis(2-nitrobenzoic acid). Analytical Biochemistry 175:408–413.

Ugalde J.M., Fecker L., Schwarzländer M., Müller-Schüssele S.J., Meyer A.J. (2022) Live monitoring of ROS-induced cytosolic redox changes with roGFP2-based sensors in plants. In: Mhamdi A (ed) Reactive Oxygen Species in Plants. Springer US, New York, NY, pp 65–85.

Ugalde J.M., Fuchs P., Nietzel T., Cutolo E.A., Homagk M., Vothknecht U.C., Holuigue L., Schwarzländer M., Müller-Schüssele S.J., Meyer A.J. (2021) Chloroplast-derived photo-oxidative stress causes changes in H2O2 and *EGSH* in other subcellular compartments. Plant Physiology 186:125–141.

Wagner S., Steinbeck J., Fuchs P., Lichtenauer S., Elsässer M., Schippers J.H.M., Nietzel T., Ruberti C., Van Aken O., Meyer A.J., Van Dongen J.T., Schmidt R.R., Schwarzländer M. (2019) Multiparametric real-time sensing of cytosolic physiology links hypoxia responses to mitochondrial electron transport. New Phytologist 224:1668–1684.

Watanabe K., Breier U., Hensel G., Kumlehn J., Schubert I., Reiss B. (2016) Stable gene replacement in barley by targeted double-strand break induction. Journal of Experimental Botany 67:1433–1445.

