## Supplemental Figures for "High robustness of cytosolic glutathione redox potential under combined salt and osmotic stress in barley as revealed by the biosensor Grx1-roGFP2"

**A**

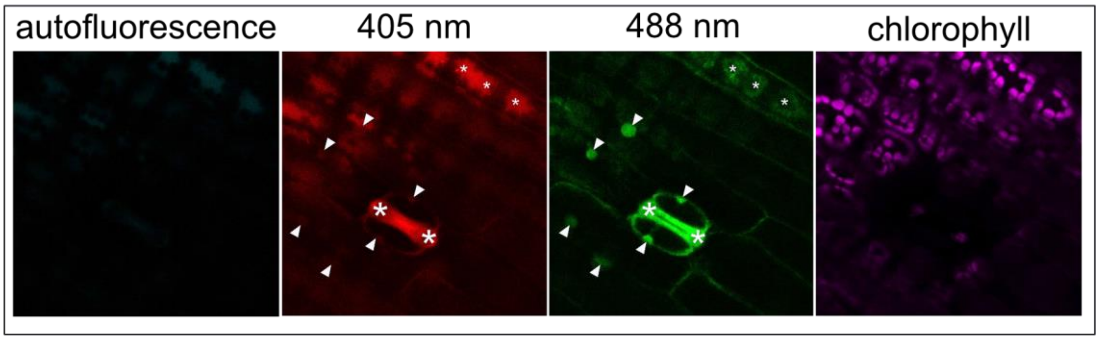


**B**

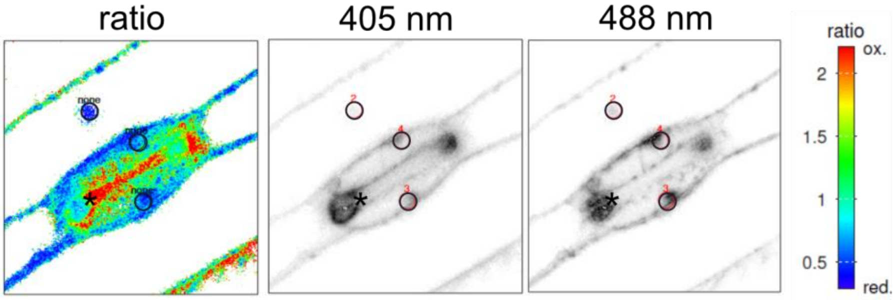


**Supplemental Figure 1: Screening and analysis of Grx1-roGFP2 signal in barley**

**A** Localisation and fluorescence signal of Grx1-roGFP2 in barley leaves. White arrowheads indicate specific roGFP2 signal, asterisks show positions of high autofluorescence signal. **B** Regions of interest (ROI, black circles) set in nuclei for ratio calculation with the RRA software in the Grx1-roGFP2 channels (405 nm, 488 nm) and the false coloured ratio image (ratio). Black asterisks show autofluorescence signal, seen as red false colouring in the ratio image.

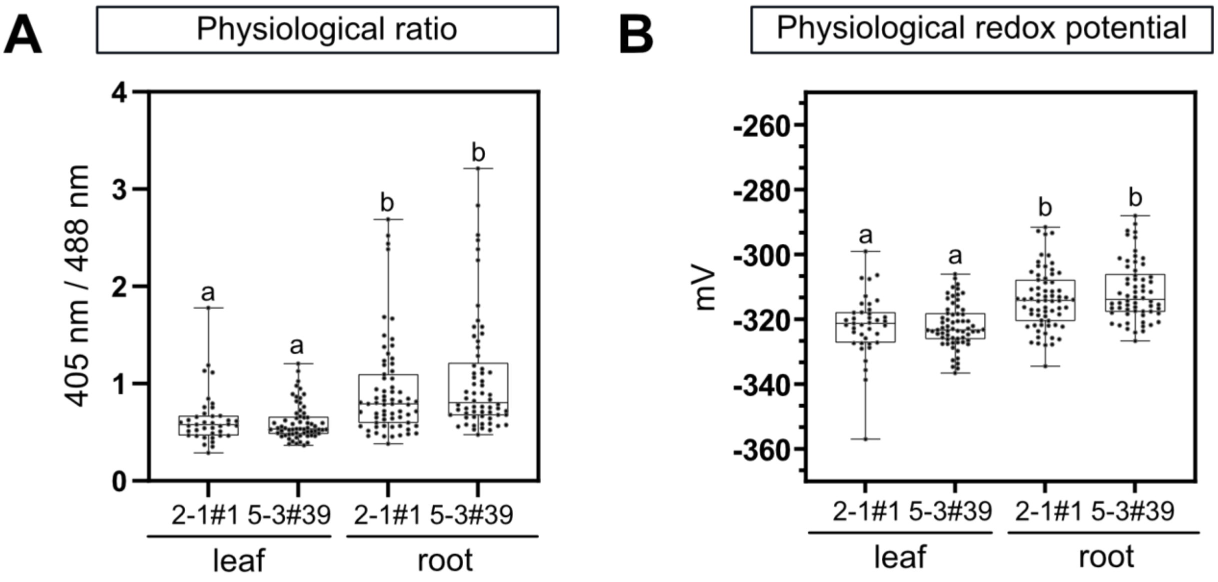


**Supplemental Figure 2: Physiological glutathione redox potential as indicated by Grx1-roGFP2 in barely root and leaf tissue**

**A** Box plots of Grx1-roGFP2 signal ratios (405/488) under physiological conditions (same data as in **Fig. 2**, but with a different scale in the y-axis to facilitate visualization). **B** The redox potential of each ratio point depicted in A was translated into mV by setting the calibration of line 2-1 #1 (DTT and DPS root values) as 100 % oxidized and reduced for a pH of 7.2 (Schwarzländer et al. 2008). Box plot features and statistics are as described in **Fig. 2 A**.

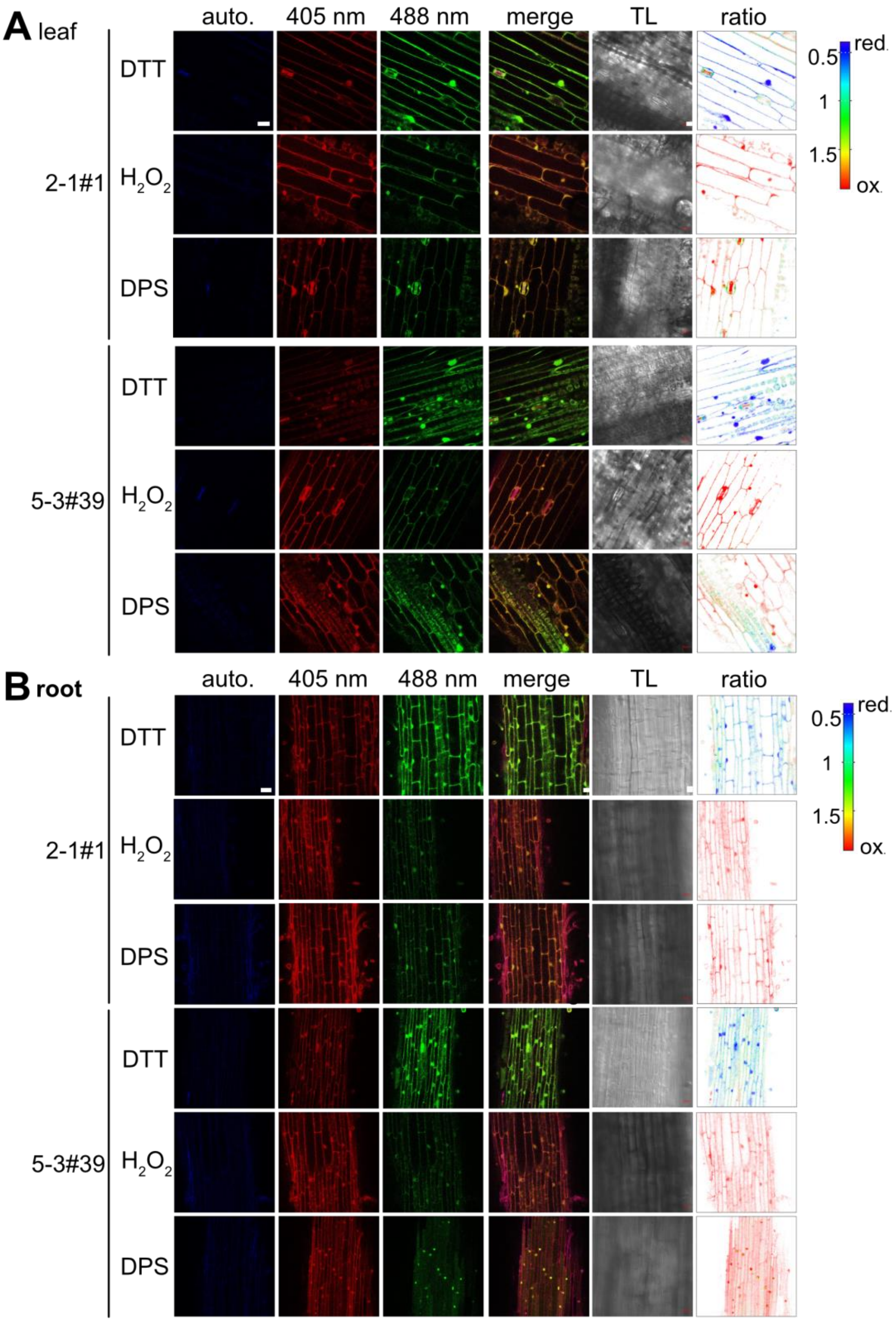


**Supplemental Figure 3: Calibration of barley root and leaf tissue**

Representative examples of the type of confocal microscopy images used to collect the data in **Fig. 2** for **A** leaf and **B** root tissues. Confocal laser scanning microscopy of barley seedlings treated with 10 mM DTT or 1.5 M $H_2O_2$ or 5 mM DPS for 30 min. roGFP2 was excited at 405 and 488 nm while emission was set to 508-535 nm. Autofluorescence was collected after excitation at 405 nm and emission between 430 to 470 nm. Overlay of channels 405 and 488 are shown as ‘merge’. Transmitted light (TL). Ratios were calculated with the ratio imaging software RRA and are false color coded. Scale bar = 40 µm.

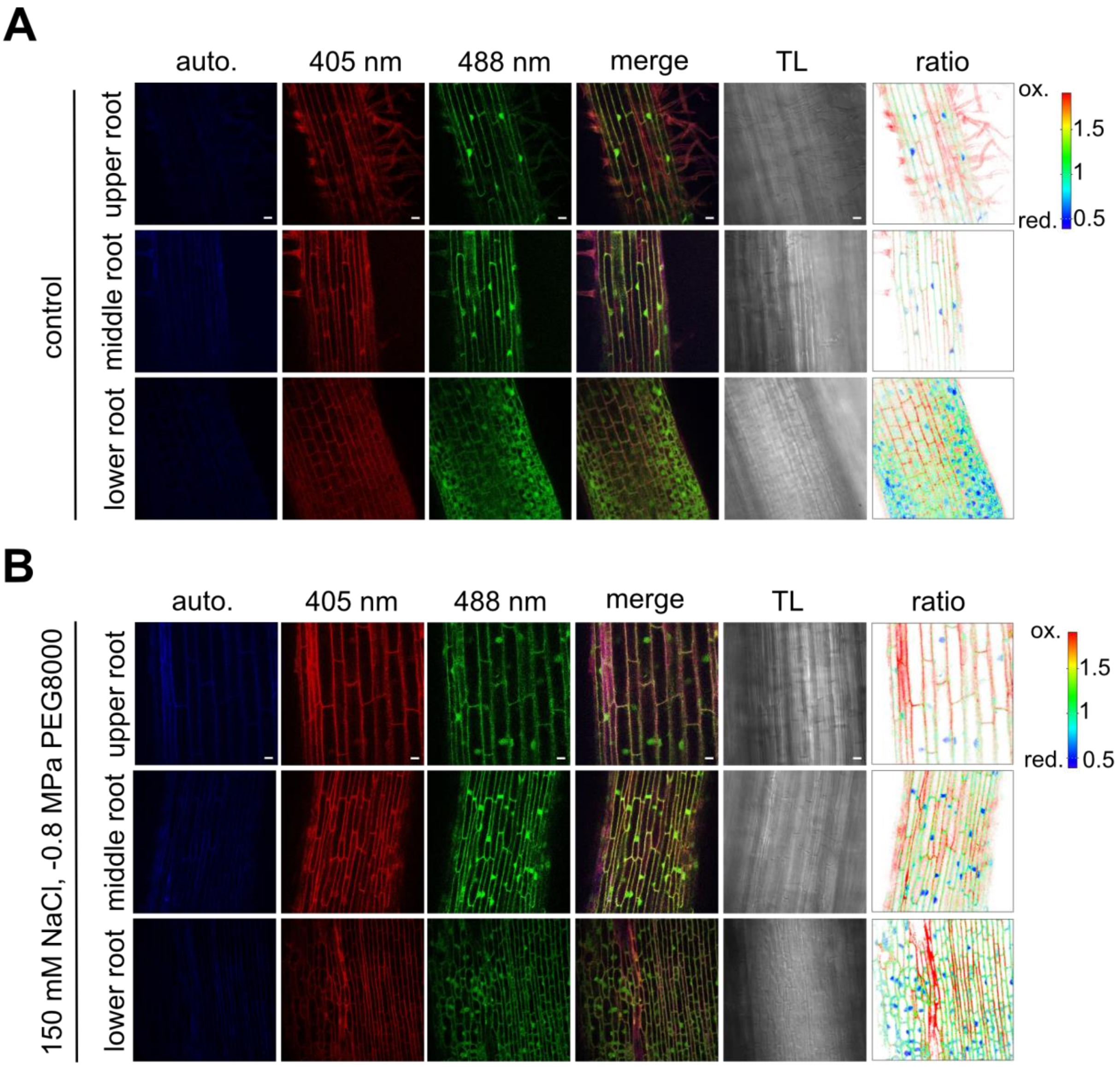


**Supplemental Figure 4: Grx1-roGFP2 ratio after 24 h of combined stress treatment**

Representative examples of the type of confocal microscopy images used to collect the data in **Fig.4** for **A** control and **B** stressed samples treated with 150 mM NaCl and -0.8 MPa PEG800 for 24 h. roGFP2 was excited at 405 and 488 nm and emission set to 508-535 nm. Autofluorescence was collected after excitation at 405 nm and emission set to 430 to 470 nm. Overlay of channels 405 and 488 are shown as ‘merge’. Transmitted light (TL). Ratios were calculated with the ratio imaging software RRA and are false color coded. Scale bar = 40 µm.

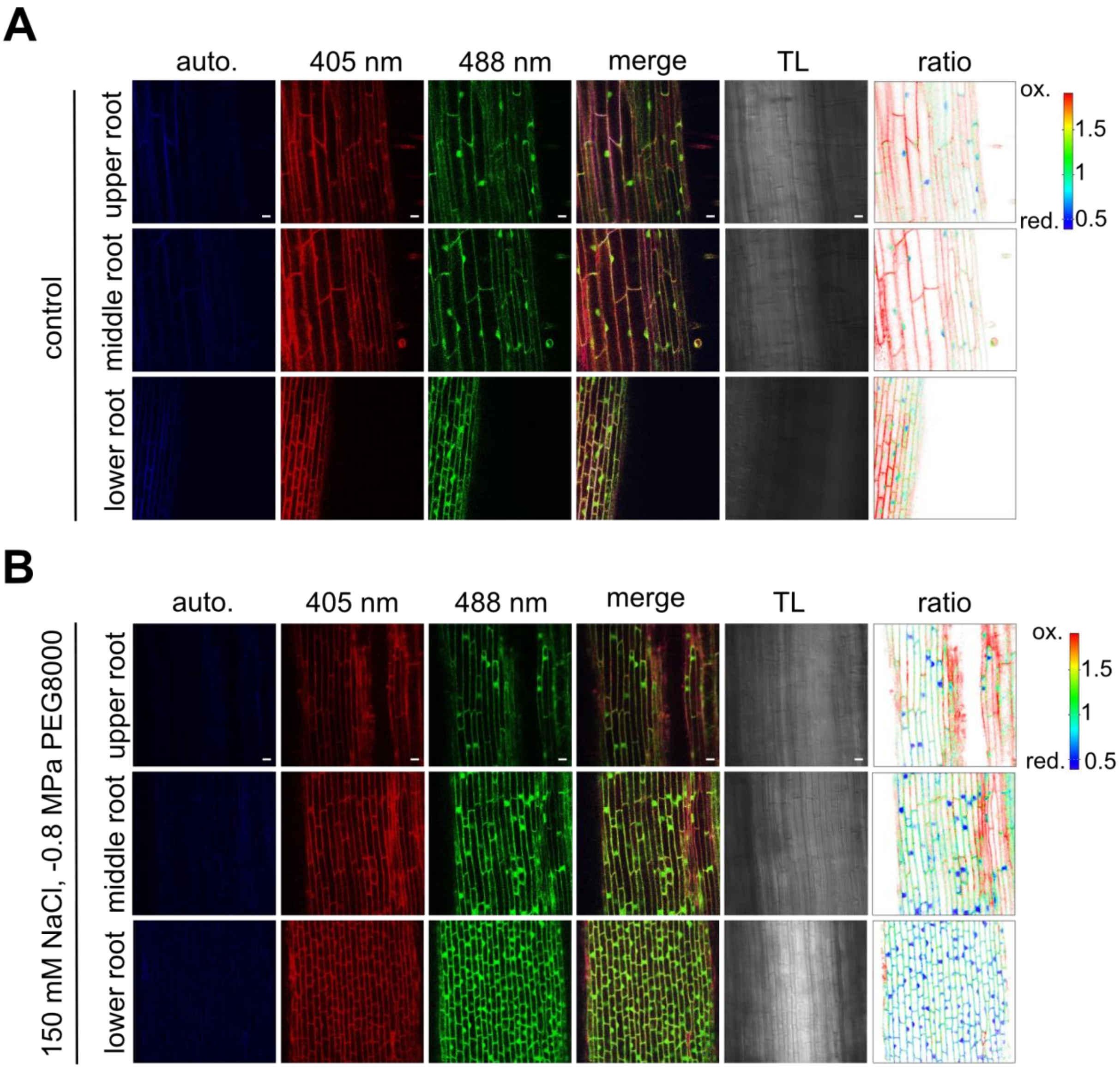


**Supplemental Figure 5: Grx1-roGFP2 ratio after 48 h of combined stress treatment**

Representative examples of the type of confocal microscopy images used to collect the data in **Fig.4** for **A** control and **B** stressed samples treated with 150 mM NaCl and -0.8 MPa PEG800 for 48 h. roGFP2 was excited at 405 and 488 nm and emission set to 508-535 nm. Autofluorescence was collected after excitation at 405 nm and emission set to 430 to 470 nm. Overlay of channels 405 and 488 are shown as 'merge'. Transmitted light (TL). Ratios were calculated with the ratio imaging software RRA and are false color coded. Scale bar = 40 µm.

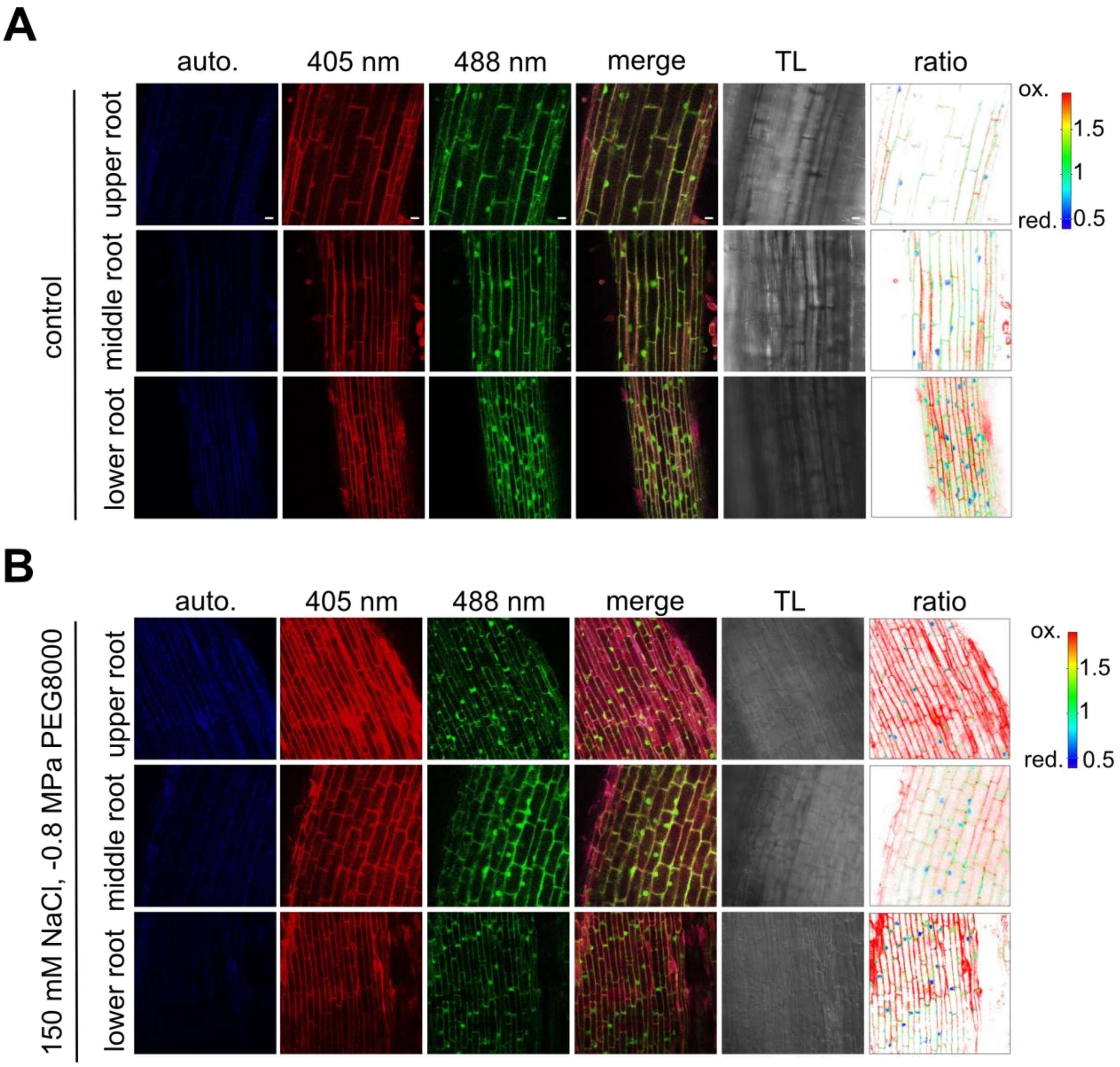


**Supplemental Figure 6: Grx1-roGFP2 ratio after 96 h of combined stress treatment**

Representative examples of the type of confocal microscopy images used to collect the data in **Fig.4** for **A** control and **B** stressed samples treated with 150 mM NaCl and -0.8 MPa PEG800 for 96 h. roGFP2 was excited at 405 and 488 nm and emission set to 508-535 nm. Autofluorescence was collected after excitation at 405 nm and emission set to 430 to 470 nm. Overlay of channels 405 and 488 are shown as 'merge'. Transmitted light (TL). Ratios were calculated with the ratio imaging software RRA and are false color coded. Scale bar = 40 µm.
